# RI-SINDy: Regulation-Informed Sparse Model Discovery in Biological Gene Networks

**DOI:** 10.64898/2026.10.04.756519

**Authors:** Nicholas J. Lucenti, Majid Bani Yaghoub

## Abstract

Oscillatory and saturating dynamics in regulatory gene networks govern processes such as cell cycle timing, circadian rhythms, and inflammatory signaling, and sparse regression methods such as SINDy offer an interpretable method for recovering their governing equations from time-series data. However, these networks are typically governed by Hill-type nonlinearities that become nearly collinear with polynomial candidates over most bounds of an oscillating curve, and placing them in a shared library produces ill-conditioned regressions and unreliable coefficient estimates. Standard approaches such as ridge regularization, library normalization, and orthogonalization can marginally improve conditioning but cannot remove the structural multicollinearity of these systems, in turn limiting our ability to produce accurate parameter estimates. To address this issue, we propose the Regulation-Informed Sparse Identification of Nonlinear Dynamics (RI-SINDy) method, which removes known regulatory terms from the regression and recovers their coefficients directly from an asymptotic drift-balance condition. Because the time averaged derivative of a bounded trajectory approaches zero, the coefficients of regulatory nonlinearities can be computed independently by constraining them to offset the mean contribution of the remaining terms. We demonstrate that this method effectively reduces library collinearity and improves conditioning numbers by 3–45*×*, substantially improving coefficient recovery and regression accuracy. We test the RI-SINDy method on experimental Hes1 data and on synthetic Goodwin and NF-*κ*B models. On Hes1, it achieves the lowest trajectory error among the compared methods (Traditional SINDy, SR3, and Nullcline Reconstruction). On the synthetic systems, it recovers the correct sparse structure with lower trajectory error than SR3 and Nullcline Reconstruction, and in a Goodwin noise sweep it recovers all active terms at noise levels up to 15%, where Traditional SINDy and E-SINDy fail to. In conclusion, the RI-SINDy method provides a robust framework to overcome the issue of multicollinearity for system identification in biological systems.

## 1 Introduction

Oscillatory and saturating dynamics in gene expression govern a variety of processes in cell biology, including cell cycle timing, developmental signaling, circadian and ultradian rhythms, and inflammatory signaling [1–7]. Mechanistic differential equation models of these processes have served as a powerful tool for relating kinetic parameters on a molecular level to population level behaviors, as well as for generating predictions about how regulatory networks will respond to perturbation [8–10]. Traditionally, the construction of these models required substantial prior mechanistic knowledge of the systems they describe, which produced large limitations in the context of largely misunderstood or unstudied gene networks.

Data-driven model discovery has emerged as an alternative approach to this problem, and in recent years we have seen a rise in algorithms such as the Sparse Identification of Nonlinear Dynamics (SINDy) framework [11]. By expressing a system’s governing equations as a sparse linear combination of candidate functions, SINDy produces models that remain mechanistically interpretable in a way that black-box machine learning approaches typically do not, and this property has made it particularly useful in cell biology, where a parsimonious closed form equation can be directly related to a meaningful hypothesis [12, 13]. When applied directly to real biological time-series, however, SINDy and many of its variants frequently struggle to recover stable, biologically plausible models, a limitation documented across many efforts to apply these methodologies to experimental data [12]. A common cause is the Hill-type and other saturating nonlinearities that govern activation, repression, and cooperative binding in regulatory and signaling networks [1, 5, 14, 15]: over the restricted trajectory a biological system actually traces out, these saturating functions can become highly collinear with the polynomial and scalar candidates already present in the library, and this collinearity has been shown to degrade the conditioning of the regression [16–18]. Because Hill-type kinetics appear throughout regulatory biology, this conditioning problem limits many existing model identification frameworks from producing meaningful insight into these systems.

The SINDy literature has approached this instability from several directions, each targeting a specific symptom of the underlying problem. Ensemble based approaches such as E-SINDy improve robustness to sampling noise by aggregating coefficient estimates across bootstrapped subsets of the data and library [19], and Bayesian formulations such as UQ-SINDy replace point estimates with full posterior distributions under sparsity promoting priors, allowing poorly supported terms to be flagged through their uncertainty distributions rather than through a hard threshold [20]. Weak-SINDy reformulates the regression in an integral rather than a pointwise sense, which improves robustness to high frequency noise [21]. These approaches improve robustness through different mechanisms, but all continue to estimate the regulatory nonlinearities’ coefficients within the shared regression itself. None of these approaches directly target the specific candidate terms responsible for most of the ill-conditioning, so the underlying collinearity between saturating terms and the surrounding library tends to persist across bootstrapping, sparsity priors, and weak formulations alike. Implicit formulations of SINDy address rational nonlinearities, of which Hill functions are a special case, by placing the derivative on both sides of the regression and identifying numerator and denominator polynomials jointly [22, 23]. Because a Hill function of cooperativity *n* has a degree-*n* polynomial denominator, the enlarged library grows with *n*, and all coefficients are still estimated in a single regression. RI-SINDy instead retains the known parametric Hill form, so its coefficient remains directly interpretable, and removes it from the regression. A separate line of work extends SINDy to libraries containing parametric candidate functions, jointly optimizing embedded nonlinear parameters alongside the linear coefficients through a nonconvex relax-and-split scheme [24]. Another recent advancement is the Nullcline reconstruction variant, which identifies inaccuracies in the inferred position of a system’s limit cycle in phase space as a separate source of instability, refitting the library after offsetting the data to better localize the system’s nullclines [25]. This correction targets the identification of stable systems effectively, but can still be affected by the same ill-conditioning in real applications. A different approach replaces the unknown nonlinearity with a neural network, fitting a hybrid mechanistic and neural model before applying a sparse regression to the network’s output [26, 27], which evades the conditioning problem since a neural network’s fit is not subject to the same collinearity constraints as a linear regression, but with a loss of interpretability, since the recovered Hill-type behavior becomes a black-box function rather than a coefficient scaling a known kinetic formulation.

This leaves a gap in our ability to make interpretable insights into systems governed by saturating kinetic processes, since existing approaches either retain the term inside the regression, where it degrades conditioning regardless of how it is regularized, repositioned, or made robust to noise, or they replace it with a function whose shape is learned rather than known. Rather than further modifying the regression itself, this work aims to supplement it with an external source of physical information. Imposing known physical structure on sparse regression has precedent in constrained SINDy [28], which enforces energy-preserving quadratic structure; here the constraint is an asymptotic balance on the mean of each state derivative that is applied to determine a single coefficient.

For a bounded oscillatory trajectory, the time average of the state derivatives must approach a zero average as the system evolves [29]. RI-SINDy exploits this balance when the contribution of a known regulatory mechanism is the remaining unknown term in an equation: the prolonged average of the other library terms’ contributions determines the regulatory contribution required to maintain zero net drift. This work introduces Regulation-Informed Sparse Identification of Nonlinear Dynamics (RI-SINDy), a framework developed specifically for system identification within bounded, oscillatory dynamical regimes, which formalizes this balance to decompose the candidate library and remove regulatory terms from the sparse regression itself. Shrinking the candidate library to a set of well-conditioned standard basis functions simplifies the regression, allowing standard sparsifying algorithms, uncertainty quantification, and noise filtering processes to operate on the remaining polynomial dynamics with improved accuracy [30–33]. The decoupled regulatory terms are then recovered by enforcing this drift-balance constraint across the limit cycle, and constraining the identification to explicitly satisfy this relationship yields models that are generally more physically interpretable, since in real biological systems, production and degradation mechanisms naturally act in opposition to maintain bounded oscillations, and embedding this opposing dynamic directly into the parameter identification prevents non-physical solutions. We demonstrate RI-SINDy on canonical synthetic genetic oscillators and experimental time-series, comparing its performance against Nullcline Reconstruction, SR3, Ensemble SINDy, and Traditional SINDy [19, 25, 34] across varying degrees of library collinearity and measurement noise.

We derive this drift-balance condition formally in §2.4. Because the drift constraint assumes a prolonged balance, the method is expected to be less reliable for trajectories that are incompletely observed, which is further discussed in §4. This proposed framework is not inherently restricted to Hill kinetics, and is realistically applicable to any regulatory nonlinearity whose mechanistic form is known but whose coefficient is difficult to identify within a shared sparse regression. We focus on Hill-type saturation here because of its prevalence in transcriptional regulation, and test it on various systems that pertain to this type of biology.

The remainder of this paper is organized as follows. §2 presents the methodology of this work. §2.1 introduces the SINDy paradigm and the notation used throughout the remainder of the paper. §2.2 introduces RI-SINDy, detailing the decomposition of the candidate library and the observed derivative data into standard basis and regulatory components. §2.3 describes the Iteratively Reweighted Least Squares procedure used for Bayesian uncertainty quantification on the standard basis regression, §2.4 derives the drift-balance condition used to identify the regulatory terms’ coefficients, and §2.5 describes the propagation of uncertainty into those coefficients via the delta method. §3 presents the application of RI-SINDy to the Hes1 protein-mRNA network, the Goodwin attractor, and the NF-*κ*B signaling network, including a direct numerical comparison against SR3, Nullcline Reconstruction SINDy, and Traditional SINDy.

## 2 Methodology

### 2.1 Sparse Biological Model Identification

The Sparse Identification of Nonlinear Dynamical Systems paradigm recovers the governing equations of a dynamical system directly from time-series data. SINDy identifies functions of the form 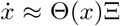, where Θ ∈ ℝ^*m×p*^ is a library of candidate terms for the regression to select from, Ξ ∈ ℝ^*p×j*^ is the coefficient matrix, and 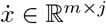 is the matrix of observed derivatives [11].

Each column of Θ corresponds to one candidate function evaluated across every sampled time point *t*_*n*_, so that the rows of Θ index time and the columns index candidate terms. A commonly used choice is the polynomial library

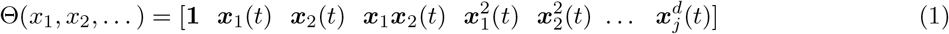

and the observed derivative data is arranged in the corresponding matrix

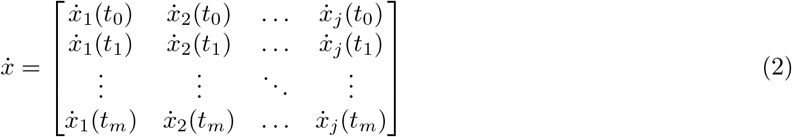

The coefficient matrix Ξ ∈ ℝ^*p×j*^ contains the coefficient vector for each dimension *j* of the system, Ξ = [***ξ***_1_ |***ξ***_2_ |… | ***ξ***_*j*_]. The derivatives in 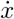 are estimated from the raw measurements using Savitzky-Golay filtering in MATLAB, which performs well under low to moderate measurement noise. The coefficient matrix is recovered by taking a pseudo-inverse of Θ and applying it to 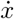, with the regression repeated iteratively and a hard threshold applied after each pass to zero out any term whose magnitude falls below the sparsity parameter *λ*.

### 2.2 Proposed RI-SINDy Method

A core difficulty in applying SINDy to regulatory gene networks is that the candidate library typically has to include a saturating, Hill-type nonlinearity alongside ordinary polynomial and scalar terms. RI-SINDy addresses this by removing the saturating term from the regression entirely and identifying its coefficient through an independent asymptotic drift-balance constraint, to improve regression conditioning and internally enforce physical properties. Figure 1 illustrates the key distinction between Standard SINDy and the proposed RI-SINDy methodology. Rather than estimating highly collinear regulatory and polynomial terms within a single sparse regression, RI-SINDy separates known regulatory terms and recovers their coefficients independently through the drift-balance condition. This decomposition yields a better-conditioned sparse regression while preserving the mechanistic interpretation of the regulatory dynamics.

**Figure 1:**
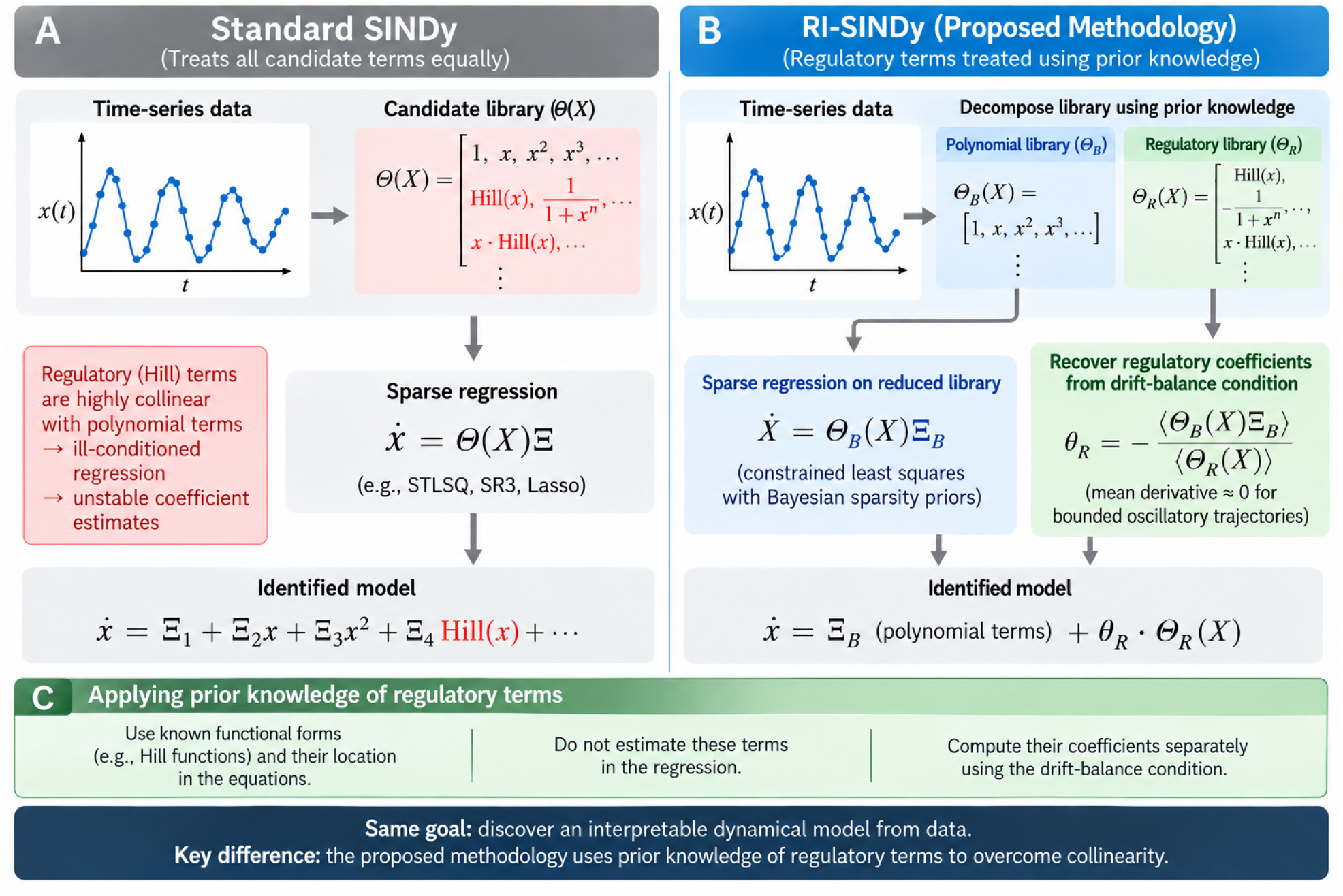
Comparison of Standard SINDy and RI-SINDy. RI-SINDy separates known regulatory terms from the standard candidate library and estimates their coefficients using a drift-balance condition to reduce collinearity and improve regression conditioning.

The RI-SINDy procedure operates by alternating between two updates until both converge on a set of coefficients for the identified system: the standard basis coefficients are fit by constrained least squares against the derivative data with the current regulatory terms’ contributions subtracted out, and the regulatory coefficients are computed from the updated standard basis coefficients using the drift-balance condition derived in §2.4. Algorithm 1 summarizes this outer procedure, with the regulatory coefficient and its uncertainty computed at each pass via the function detailed in Algorithm 2.

We split the candidate library into an augmented matrix Θ = [Θ_*ℬ*_|Θ_*ℛ*_],

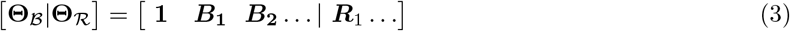

where **Θ**_*B*_ contains the standard basis terms: the polynomial, scalar, and signal candidates the pseudo inverse identifies without difficulty. **Θ**_*R*_ contains the regulatory terms: the saturating Hill-type functions describing activation, repression, or other switch-like regulatory behavior in the network. We refer to the coefficients of **Θ**_*R*_ as the regulatory coefficients, since they are identified not through the shared sparse regression but through the procedure in §2.4.

The regression begins with a traditional least squares estimate over the full library,

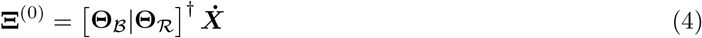

This estimate only sets up the first drift-balance update; Ξ_*B*_ is refit on the reduced library for all future passes. Since the system’s derivative state is the sum of the contributions from the standard basis terms and the regulatory terms,

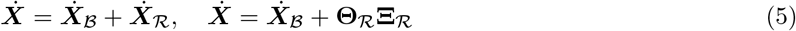

Once the regulatory contribution ***I*** = **Θ**_*R*_**Ξ**_*R*_ has been estimated, it is subtracted from the observed derivative data,

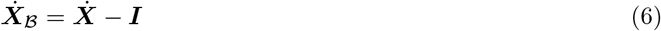

which leaves a derivative target that the standard basis terms must independently explain. We use the notation ***I*** for this influence term to keep the algorithm readable, since it is the quantity subtracted out before each pass of the standard basis regression. With the regulatory term removed, the standard basis regression resolves only the well conditioned part of the library, which is expected to improve identification of the active terms. The standard basis coefficients are found via constrained least squares (LSQLIN). Each pass of the standard basis regression solves

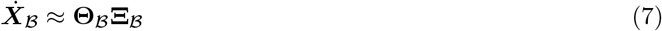

The alternating procedure can be interpreted as a block-coordinate estimation: one block consists of a constrained regression over the standard-basis coefficients, while the second block is obtained from the closed-form drift-balance constraint. The regulatory update does not minimize the same objective as the least-squares subproblem; instead, it enforces the trajectory balance condition. In practice, relaxation of both updates, via the parameter *η* introduced in §2.3 and §2.5, helps produce stable convergence.

The conditioning benefit of this decomposition follows from removing the regulatory terms from the matrix that is inverted. Over the range of a bounded trajectory, a Hill column can be closely approximated by combinations of polynomial columns, making the full library nearly rank deficient. Library columns are normalized to unit *l*_2_ norm before regression and coefficients are rescaled afterward; condition numbers are computed on the normalized library, so differences in column scale do not contribute. Figure 2(c) shows, for a synthetic Goodwin model, that appending Hill columns to a polynomial library of degree *d* increases the condition number by a factor *κ/κ*_*B*_ of 38.9, 42.0 and 3.8 for *d* = 1, 2, 3 with two or more Hill terms. We also quantify the conditioning reduction of mean centered and variance normalized libraries Figure 2(a) to assess the impact of Hill decoupling in the context of decorrelated data, where we find only marginal improvements over the raw data. Figure 2(b) shows the application of a ridge regularization on the collinear library, which displays slight improvements in highly correlated libraries but retains the same flaw of struggling to scale with the worsening condition in libraries with multiple Hill functions.

**Figure 2:**
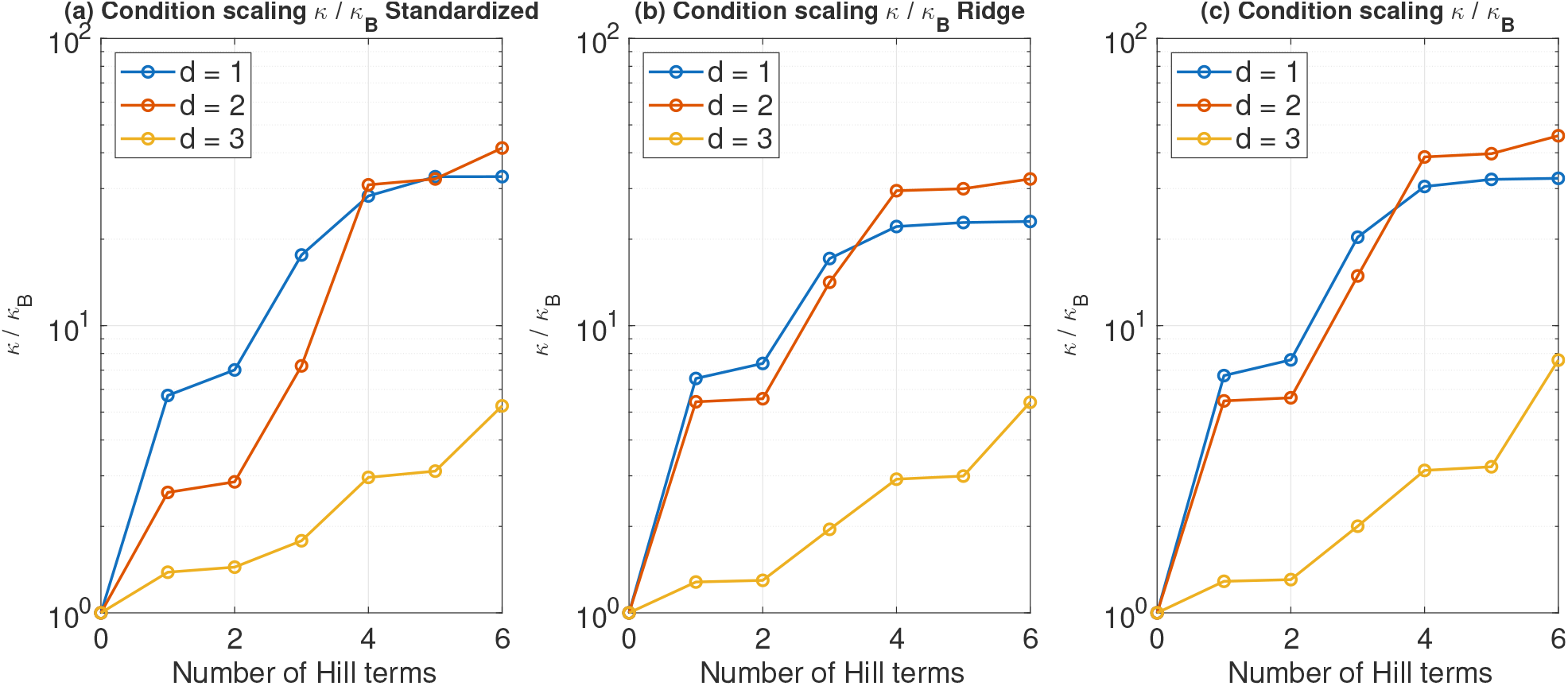
Effect of Hill terms on library conditioning, using Goodwin oscillator limit-cycle data (*n* = 10). Ratio of the condition number of the standardized (mean and variance normalized) library to that of the polynomial only library of the same degree *d*, as Hill terms are added. (b) The same ratio under ridge regularization. (c) Condition ratio of the raw library.

### 2.3 Uncertainty Quantification

Sparsity inducing priors are built into this framework so that approximate posterior distributions can be identified alongside the coefficients themselves, placing more probability mass near zero for terms that end up thresholded out while leaving active terms largely unpenalized [30]. The Horseshoe prior [35] is well suited to regulatory network models, where most candidate terms’ coefficients are expected to be either negligibly small or entirely absent. The priors are implemented through Iteratively Reweighted Least Squares [20].

#### Algorithm 1

RI-SINDy: sparse regression with regulatory decoupling

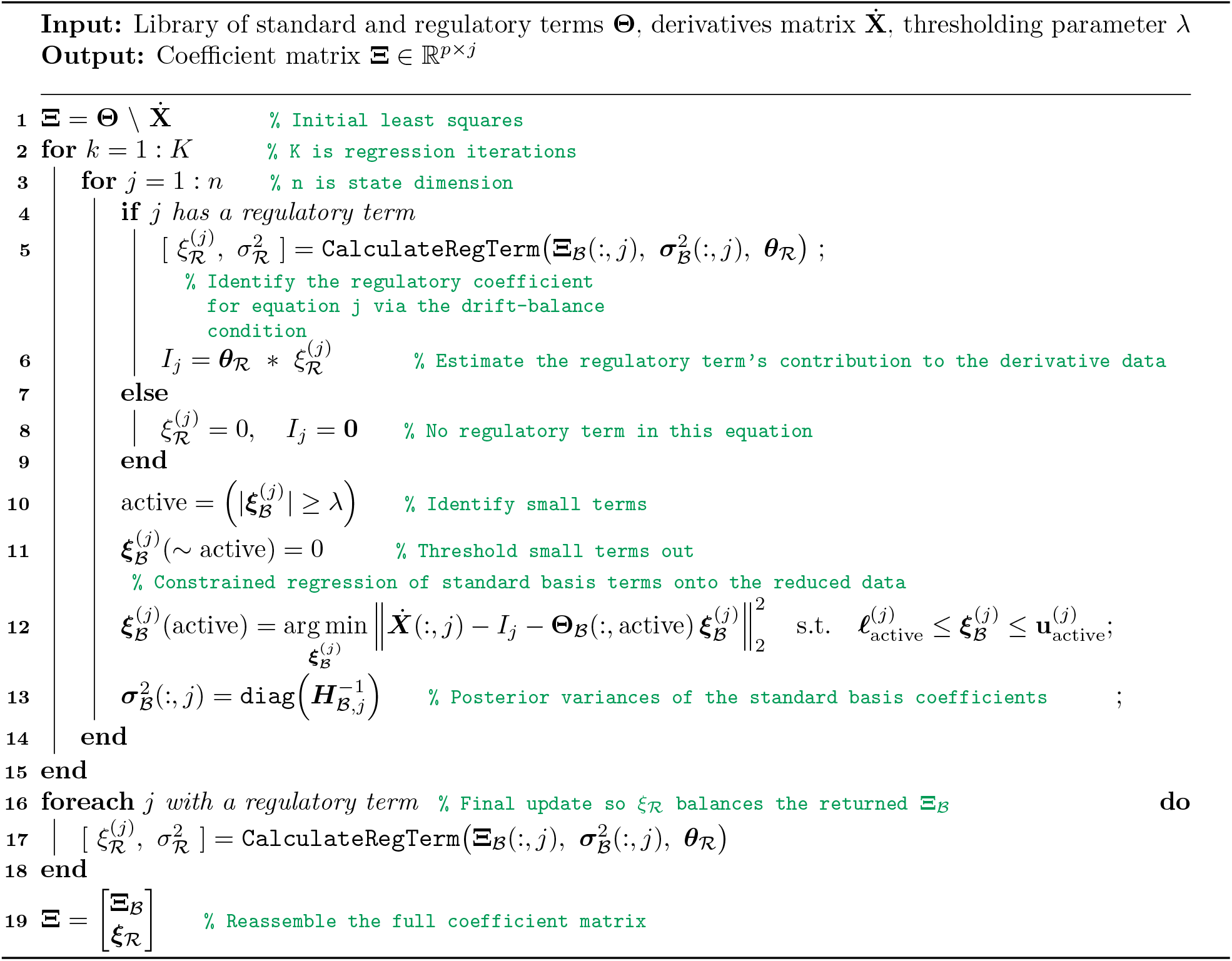

The uncertainty quantification in RI-SINDy differs from applying the same Bayesian framework directly to Standard SINDy. In Standard SINDy, both standard-basis and regulatory coefficients are estimated jointly from the full candidate library, so collinearity between polynomial and regulatory terms is inherited by the posterior coefficient estimates. In RI-SINDy, Bayesian uncertainty quantification is instead performed on the reduced standard-basis library Θ_B_, while uncertainty in the regulatory coefficients is subsequently propagated through the drift-balance relation derived in subsection 2.4. This can be expressed directly through the corresponding regression formulations. For Standard SINDy with UQ, the full library enters a single regression,

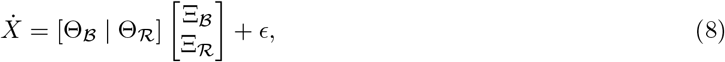

so that uncertainty in both coefficient blocks is inferred from the same potentially ill-conditioned library. In contrast, RI-SINDy performs the Bayesian regression only on the reduced standard-basis library,

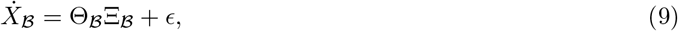

while the regulatory coefficient is determined separately through the drift-balance relation

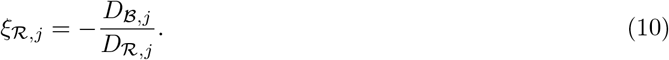

Consequently, the regulatory–polynomial collinearity is removed from the regression used to estimate Ξ_*B*_, and uncertainty in *ξ*_*R,j*_ is subsequently propagated from the standard-basis coefficients as described in subsection 2.5. The resulting distinction between the two UQ formulations is summarized in Table 1.

**Table 1:** Comparison of uncertainty quantification in Standard SINDy and RI-SINDy.

| Feature | Standard UQ SINDy | UQ RI-SINDy |
| --- | --- | --- |
| Regression library | $[\Theta_{\mathcal{B}} \mid \Theta_{\mathcal{R}}]$ | $\Theta_{\mathcal{B}}$ |
| Standard coefficients $\Xi_{\mathcal{B}}$ | Jointly estimated | Bayesian sparse regression |
| Regulatory coefficients $\Xi_{\mathcal{R}}$ | Jointly estimated | Drift-balance constraint |
| UQ of $\Xi_{\mathcal{B}}$ | Posterior from full library | Posterior from reduced library |
| UQ of $\Xi_{\mathcal{R}}$ | Posterior from full regression | Propagated from $\Xi_{\mathcal{B}}$ via delta method |
| Regulatory-term collinearity | Remains in posterior estimation | Removed from regression |

We now describe the Bayesian estimation of the standard-basis coefficients Ξ _*B*_ used in RI-SINDy. Sparsity-promoting priors are incorporated through Iteratively Reweighted Least Squares (IRLS) [35], with the prior determining the coefficient-specific weights applied during the regression. We use the Horseshoe prior as the primary formulation because it strongly shrinks unsupported candidate terms toward zero while allowing sufficiently supported coefficients to remain active [35]. Under this version of the Horseshoe prior, the weight update is

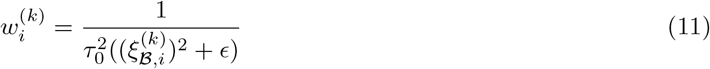

where *τ*_0_ is a global shrinkage factor and *ϵ* is a small stability constant that keeps the weight from diverging to infinity once a coefficient has been thresholded to zero. RI-SINDy can also be run with the Laplace or Spike-and-Slab priors [32, 33], which is useful for cross validating an identified term’s uncertainty under various sparsity assumptions. Their weight updates are

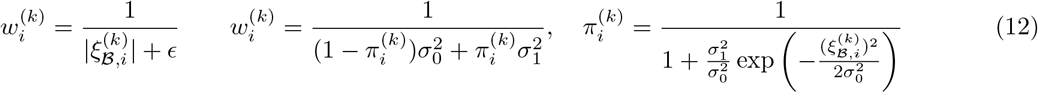

where 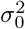 and 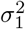 are the spike and slab variance hyperparameters. The IRLS regression operates only on the standard basis terms and data, **Θ**_*B*_ and 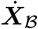, recovering only **Ξ**_*B*_. This is a weighted least squares problem in which the inverse weight matrix rescales the columns of **Θ**_*B*_,

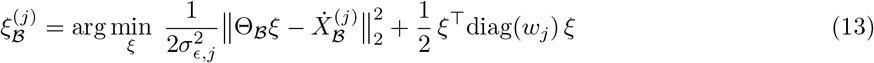

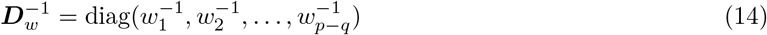

Each iteration of this reweighted regression is equivalent to a MAP estimate under the chosen sparsity prior, with the posterior covariance approximated by a Laplace expansion around that estimate [20, 36],

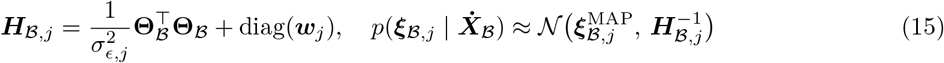

with posterior covariance 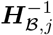, whose diagonal entries are the coefficient variances used below,

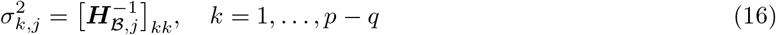

These variances feed back into the prior weights for the next iteration of the regression, and propagate into the regulatory coefficient’s variance through the delta method, described fully in §2.5. Once the variance for each term in dimension *j* has been computed, the weights *w*_*j*_ are updated with a relaxation factor *η* ∈ [0, 1] that prevents premature convergence,

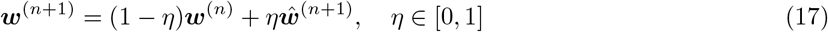

where ***ŵ***^(*n*+1)^ is the next iteration of the weight vector before relaxation. The LSQLIN regression for the standard basis terms is also subject to linear bounds 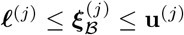, enforcing known kinetic constraints. For example, a production term can be bounded to [0, +*∞*), a degradation term to (*−∞*, 0], and an ambiguous term such as basal drift can be left open on both ends.

### 2.4 Drift Balance Identification of Regulatory Coefficients

The derivation for this constraint regards systems governed by stable oscillatory dynamics that satisfy the following three conditions on the trajectory of each state variable *x*_*j*_ used to compute the balance.

Cond 1. The trajectory must remain bounded, such that the total displacement *x*_*j*_(*T*) *− x*_*j*_(0) stays bounded as the observation window length of the data *T* grows. This is always true for a stable limit cycle, a trajectory settling toward a fixed point, or a bounded transient that decays over the window, but fails for any trajectory that accumulates without bound, such as sustained linear growth. It is worth noting here that this methodology is not applicable in the case of a monotonically growing trajectory that approaches an equilibrium that the observation window does not reach, even if an asymptotic boundedness exists, since the finite-window approximation will not encompass this convergence.

Cond 2. The discrete samples must be of uniform spacing and have a mean that adequately approximates the continuous time average over the observation window. For oscillatory trajectories, the required window depends on various factors such as oscillation period, sampling density, initial phase, transient amplitude, etc. The window enters the estimate through the boundary term [*x*_*j*_(*T*) *− x*_*j*_(0)]*/T* in (19), which decays as 1*/T* and vanishes for a periodic orbit when *T* is a whole number of periods (derived after 19). In the Goodwin test used here to quantify the effects of shifting the sampling window, the bias in *ξ*_*R*_ is below 10% for all start phases once *T ≥* 2 periods (Fig. 3).

**Figure 3:**
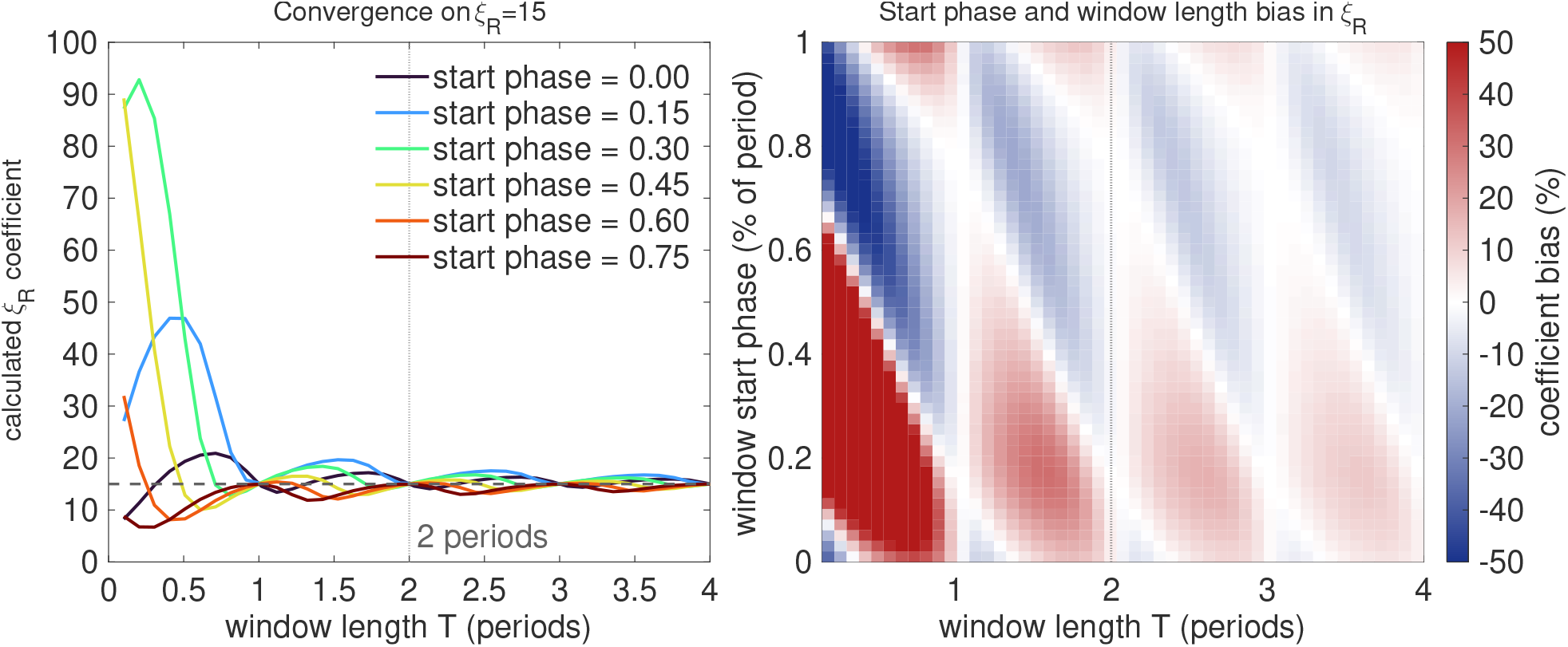
Effect of observation window length on drift-balance estimation of *ξ*_*R*_ for a Goodwin oscillator. Left: estimated *ξ*_*R*_ versus window length *T* (in periods) for six start phases, converging to the true value *ξ*_*R*_ = 15.0 (dashed) once *T* exceeds roughly two periods. Right: percent bias in *ξ*_*R*_ as a function of window length and start phase.

Cond 3. The regulatory term’s own time averaged contribution *D*_*R,j*_ must be bounded away from zero, since the recovered coefficient is obtained by dividing by it, and a regulatory term whose mean over the trajectory is near zero yields an ill-conditioned estimate regardless of how well the remaining terms are identified, a condition examined further in Figure 3.

For each equation to which the drift-balance estimation is applied, the regulatory component is represented by one known function Θ_*R,j*_(**x**) multiplied by one unknown coefficient *ξ*_*R,j*_. Under these conditions, the zero-drift constraint provides one scalar equation for one unknown regulatory coefficient. If multiple regulatory coefficients are simultaneously unknown in the same equation, the drift-balance condition alone provides only one scalar constraint and therefore does not uniquely determine all regulatory coefficients. This limitation is discussed further in Section 4.

The drift balance estimation contains three distinct steps. First, boundedness gives the asymptotic identity 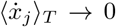 as *T→ ∞*. Second, finite observations are used to approximate this continuous time average with a discrete sample average, introducing finite-window and sampling error. Third, under the assumption that there is one regulatory term per state variable, the resulting scalar balance equation can be solved for the unknown regulatory coefficient.

We start by asserting that the time averaged derivative of a bounded state variable *x*_*j*_(*t*) converges to 0,

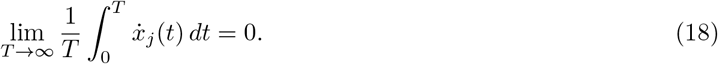

We use this asymptotic result as the basis for a discrete, finite window approximation applied to the sampled data, which is developed below.

Consider a single state variable *x*_*j*_(*t*) that does not grow or decay without bound. Its average rate of change over a window of length *T* is

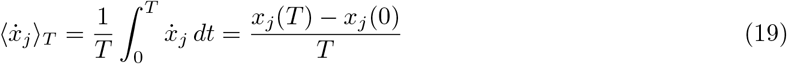

and because *x*_*j*_ stays bounded, this formulation approaches zero as *T* grows, regardless of the system’s behavior in between. We refer to 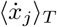 as the drift of *x*_*j*_ over the window.

Writing the identified vector field as *V* (**z**) = **Θ**(**z**)**Ξ** and substituting into the drift definition gives

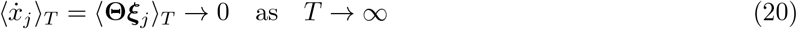

Since **ΘΞ** is a sum of term-wise contributions to 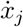, with some terms pushing the system away from its current state and others pulling it back, and since averaging is a linear operator, the average derivative 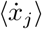 equals the sum of the average contributions from every active library term. The drift of *x*_*j*_ therefore separates cleanly into the drift contributed by the standard basis terms and the drift contributed by the regulatory term. This can be formulated as

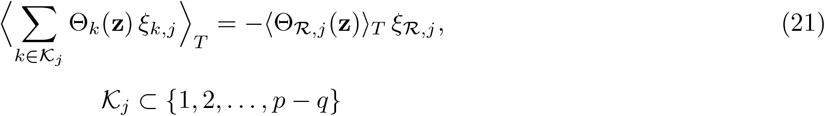

where *q* is the number of regulatory terms and *K*_*j*_ indexes the standard basis terms that are identified as active in equation *j*. Equation (21) states that the drift contributed by the standard basis terms must be matched in magnitude by the drift contributed by the regulatory term, since together they average to zero. We can thus replace the continuous time average with its discrete counterpart, evaluated directly at the sampled data. For the standard basis side this is

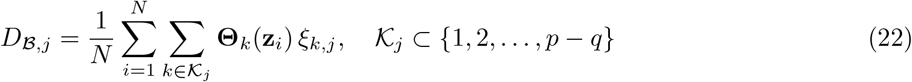

where **z**_*i*_ is the state vector (and, for explicitly time dependent terms such as *S*(*t*), the time *t*_*i*_) at the *i*th sampled time point. This evaluates every active standard basis term at each observed data point and averages the result over the full time-series. The regulatory term’s drift follows the same formulation,

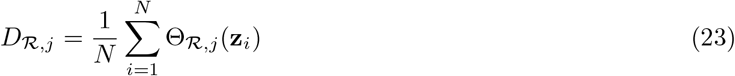

and combining the two gives the discrete drift balance condition

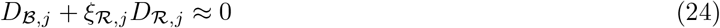

The regulatory term’s drift, scaled by its coefficient *ξ*_*R,j*_, cancels the drift produced by the standard basis terms, which is the condition a bounded trajectory must satisfy. Solving (24) for the coefficient gives

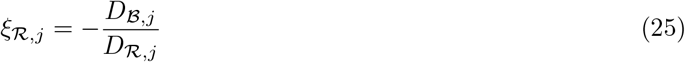

Because the active support *K*_*j*_ has already been fixed by the standard basis regression at each iteration, this is a single scalar equation in the single unknown *ξ*_*R,j*_, which yields a unique solution whenever *D*_*R,j*_ *≠* 0. This uniqueness holds regardless of the magnitude of *D*_*R,j*_, but the numerical sensitivity of the estimate does not. Because *ξ*_*R,j*_ is obtained by dividing by *D*_*R,j*_, any error in *D*_*B,j*_, whether from measurement noise, finite sampling, or an imperfectly estimated standard basis coefficient, is amplified by a factor of roughly 1*/D*_*R,j*_ in the recovered coefficient. This is the ordinary sensitivity of any division based estimator to a small denominator, and it means the drift-balance condition is best suited to regulatory terms whose own time averaged contribution to the dynamics is not itself close to zero relative to the standard basis terms it must balance against. For the positive Hill functions considered in the present experiments, *D*_*R,j*_ remains nonzero over the observed trajectories. Nevertheless, a small regulatory mean remains a potential failure mode of the estimator and should be assessed explicitly for each application.

The sign of *ξ*_*R,j*_ is determined directly by the drift-balance relation, giving *ξ*_*R,j*_ = *− D*_*B,j*_*/D*_*R,j*_, where the resulting coefficient may be positive or negative depending on the direction of the regulatory contribution, allowing the same formulation to represent activation or repression.

### 2.5 Regulatory Coefficient Variance and Update

Due to the regulatory coefficient *ξ*_*R,j*_ not being found directly by the sparse regression, its uncertainty must be estimated from the standard basis coefficients that determine it. We use the delta method [37], propagating the posterior variances of the standard basis terms through the drift-balance relation *ξ*_*R,j*_ = *−D*_*B,j*_*/D*_*R,j*_ derived in §2.4. Treating *D*_*R,j*_ as fixed and the standard basis coefficients as approximately independent, consistent with the diagonal posterior variances already reported in §2.3, the variance of *ξ*_*R,j*_ is a weighted sum of the posterior variances of the standard basis terms that are used to compute *D*_*B,j*_,

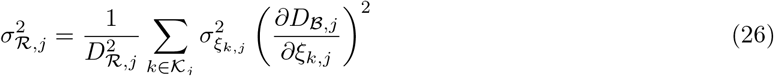

where *K*_*j*_ is again the set of active standard basis terms in equation *j*. The partial derivatives are the sample mean of each basis function over the data,

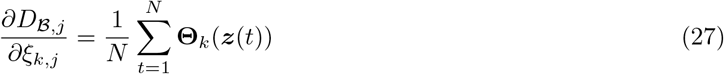

such that the uncertainty assigned to the regulatory coefficient is tied directly to the uncertainty already present in the terms that are part of the sparse regression, even though *ξ*_*R,j*_ itself is not regressed. After *ξ*_*R,j*_ is recomputed at each pass of the algorithm, it is updated with a relaxation factor *η ∈* (0, 1),

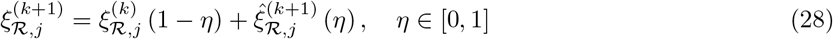

where 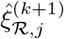 is the next iteration of this computed value before relaxation, functioning the same as the relaxation in the IRLS weight update. The full coefficient vector for equation *j* is reassembled by reconnecting the standard basis and regulatory coefficients,

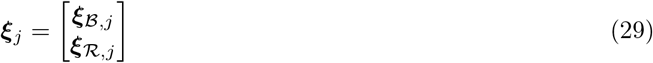

and the full coefficient matrix is reconstructed across all *j* equations,

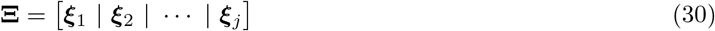

## 3 Results

### 3.1 Comparative Framework

RI-SINDy is evaluated against SR3, a parametric library extension of SINDy that optimizes an embedded nonlinear hyperparameter alongside linear coefficients [24], against Nullcline Reconstruction SINDy, a phase space offset correction method [25], and against Traditional SINDy as a benchmark. SR3 and Nullcline-SINDy are the most relevant approaches to the conditioning problems RI-SINDy attempts to resolve, which makes them the most applicable frameworks for these problems. A separate noise sensitivity sweep against Ensemble-SINDy and Traditional SINDy is reported in §3.3.2 to evaluate term identification and error robustness across noise levels.

RI-SINDy and Nullcline-SINDy are both given the true or inferred Hill hyperparameters, since neither method identifies an embedded nonlinear parameter directly; SR3 is instead given a generic initial guess, since jointly discovering these parameters is the basis for testing it. Each method’s regulatory term is also exempted from the sparsity threshold, which guarantees its activation regardless of how the sparse regression handles the remaining library terms and removes any reduced-library advantage RI-SINDy would otherwise hold.

#### Algorithm 2

Regulatory coefficient and variance evaluation

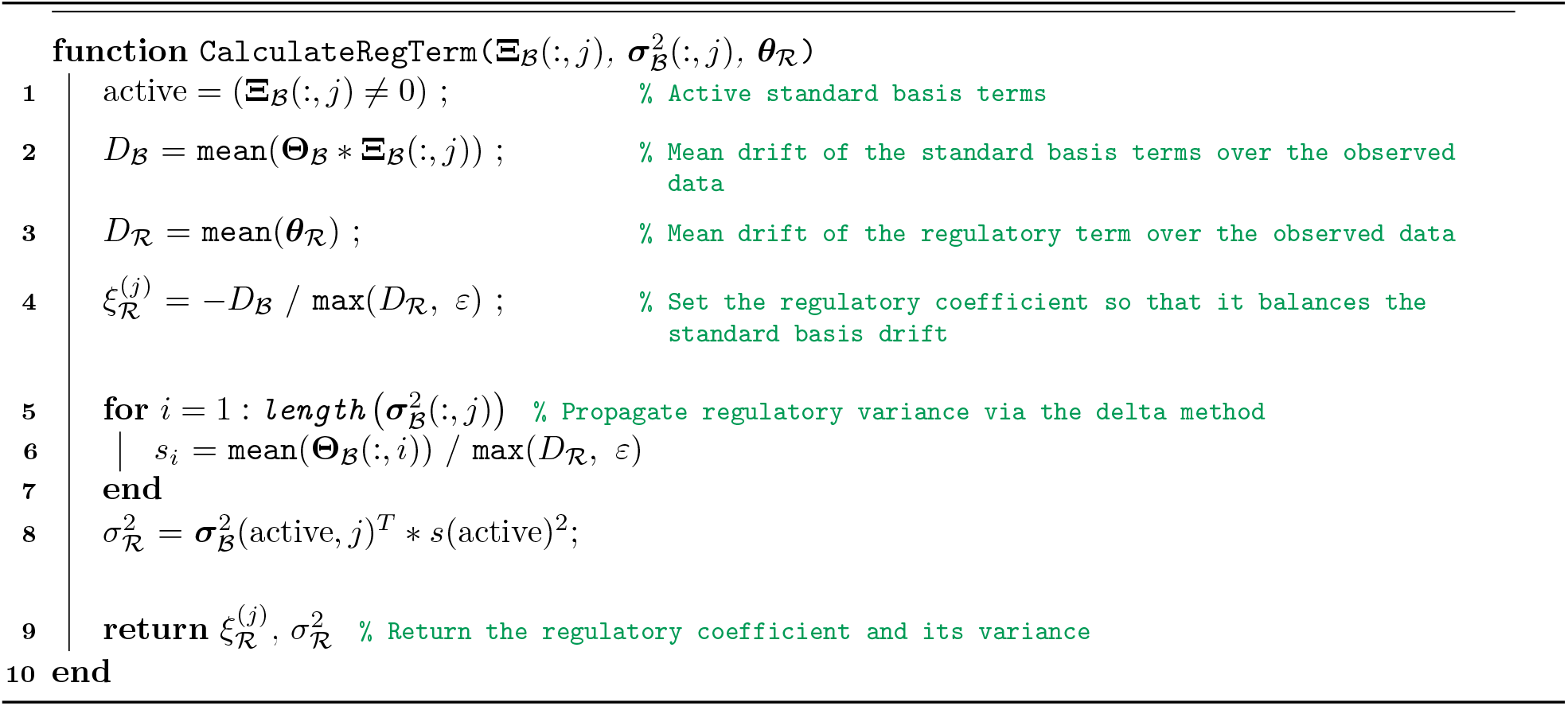

### 3.2 Hes1 Protein-mRNA Network

Hes1 (Hairy and Enhancer of Split-1) is a bHLH transcription factor involved in cell-fate regulation whose negative autoregulatory feedback can generate sustained oscillatory dynamics. Mathematical models have shown that transcriptional/translational delays can induce stable periodic solutions through Hopf bifurcation [9, 38, 39]. The Hes1 protein is an auto-repressor that inhibits the transcription of its own mRNA, and this occurs on a lagged basis which produces the observed oscillations in the Hes1 mRNA and protein concentration [1–3, 9]. The prolonged oscillatory behavior of the Hes1 system is also a product of the rapid turnover of the protein and mRNA, both of which have a half life on the order of 20 minutes [40]. The simplified dynamics for this network can be captured in the two dimensional system

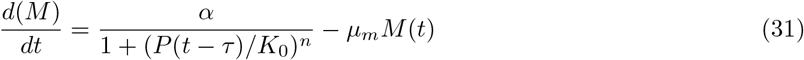

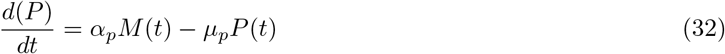

with *τ ≈* 0.3 hours. The Hill function’s coefficient *α* is identified via the drift balance condition to reduce the sparse regression to an exclusively polynomial and scalar operation. A sum of sines interpolation was applied before fitting, since the Hes1 dataset sourced from [9] contains approximately 30 measurements over a 12 hour observation period, which is insufficient for direct numerical differentiation.

The Hill hyperparameters used for Hes1, *K*_0_ = 2.7 and *n* = 9, were selected from ranges detailed in Hes1 literature [1]. Figure 4 shows the resulting pairwise correlation structure of the candidate library alongside the identified trajectories against both the interpolated and raw experimental data.

**Figure 4:**
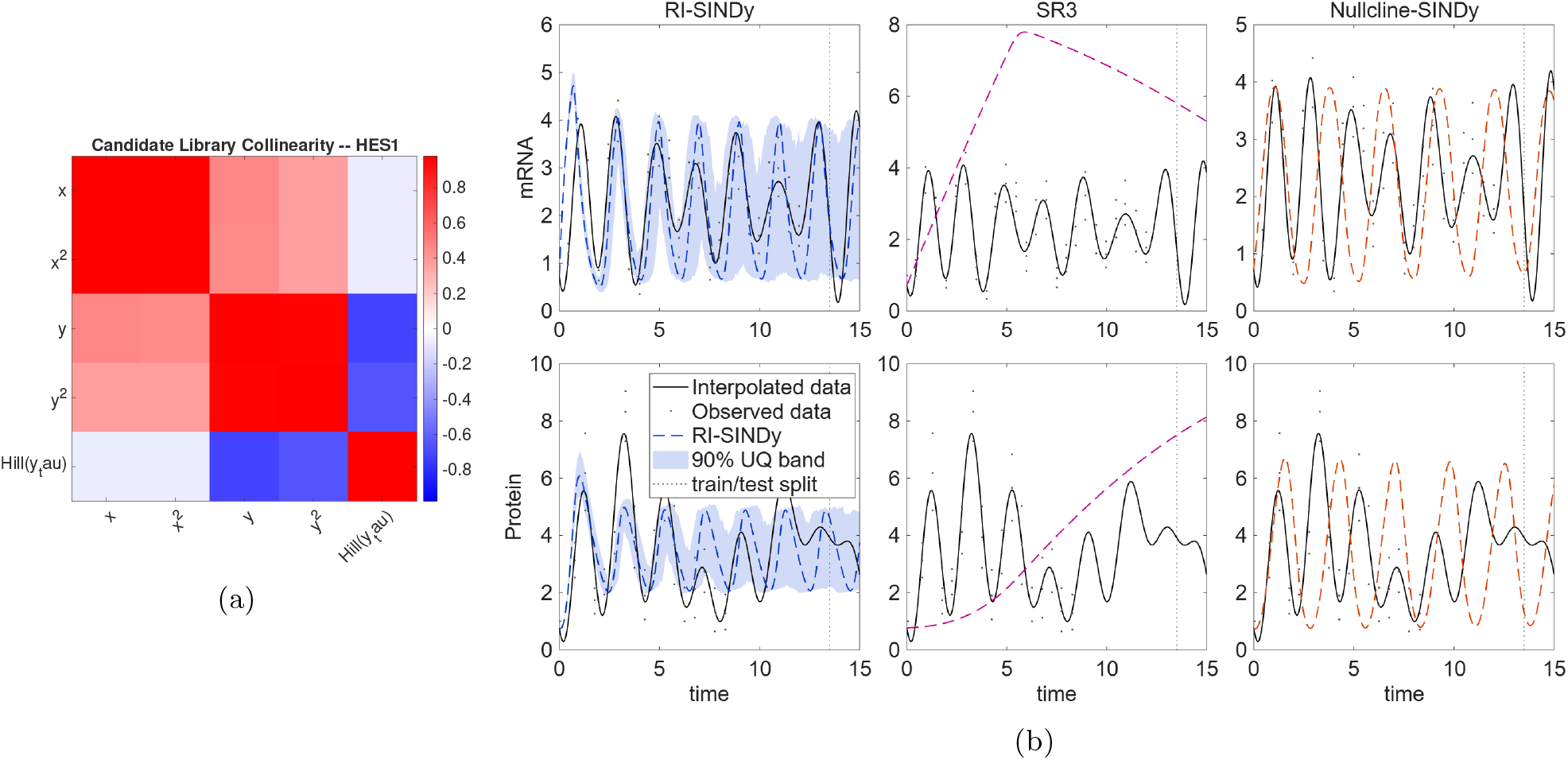
Candidate Library Collinearity and System Trajectory for the Hes1 System. (a) Pairwise correlation of all candidate library terms for the Hes1 system, evaluated on the same training data the methods are fit on. Identified trajectories against the interpolated data (black) and the observed experimental measurements (gray points). RI-SINDy’s column includes its 90% uncertainty range via Monte Carlo propagation.

The delayed Hill term is uncorrelated with either mRNA term but strongly negatively correlated with both protein terms, since this function takes a delayed state of protein concentration as an input (Figure 4a). RI-SINDy recovers a sparse two term structure in each equation: mRNA driven by linear decay and delayed Hill repression, protein driven by an *M* ^2^ production term and linear decay (Table 2). The nonzero *M* ^2^ term, rather than a simple linear production term, is potentially explained with several sources of unmodeled nonlinearity in the protein equation, one candidate being the oscillatory phosphorylation of the Stat3 protein, which has been reported to modulate Hes1 protein degradation [41, 42]. Because Stat3 oscillates in phase with Hes1 protein, such a mechanism would produce a nonlinear net accumulation of protein per unit mRNA over each period, which is the qualitative pattern RI-SINDy recovers in the *M* ^2^ coefficient without any explicit Stat3 observations in the training data. We do not have direct evidence that Stat3 is the specific mechanism responsible, and note this only as a plausible biological interpretation consistent with the recovered term rather than a confirmed source of the observed nonlinearity. SR3 assigns small nonzero weight to nearly every candidate term in both equations, while Nullcline-SINDy and Traditional SINDy return large protein equation coefficients of mixed sign (7.4671 and *−* 10.2685).

**Table 2:** Identified coefficients for the Hes1 system. RI-SINDy, Nullcline-SINDy and Traditional SINDy were given the Hill hyperparameters *K*_0_ = 2.7, *n* = 9, selected from the Hes1 literature. SR3 discovered *K*_0_ = 2.41, *n* = 20.00 (hit its upper search bound).

| Term | RI-SINDy |  | SR3 |  | Nullcline-SINDy |  | Trad. SINDy |  |
| --- | --- | --- | --- | --- | --- | --- | --- | --- |
|  | *mRNA* | *Protein* | *mRNA* | *Protein* | *mRNA* | *Protein* | *mRNA* | *Protein* |
| Constant | 0.0000 | 0.0000 | -0.0153 | 0.0053 | 0.0000 | 7.4671 | -1.3117 | -10.2685 |
| $M$ | -1.9051 | 0.0000 | -0.0056 | 0.0207 | 0.0000 | 1.0651 | -0.1721 | 1.9821 |
| $M^2$ | 0.0000 | 0.6406 | -0.0016 | 0.0098 | 0.1639 | 0.7366 | 0.0978 | 0.1096 |
| $P$ | 0.0000 | -1.0196 | -0.0104 | 0.0003 | 0.1666 | -5.6466 | 0.5445 | 0.6378 |
| $P^2$ | 0.0000 | 0.0000 | -0.0027 | -0.0001 | -0.1121 | 0.4216 | -0.1380 | 0.0000 |
| Delayed Hill | 11.7628 | 0.0000 | 1.3730 | 0.0000 | 3.8900 | 0.0000 | 3.1920 | 7.3154 |

RI-SINDy achieves the lowest trajectory error on both variables (Table 3), and its Monte Carlo uncertainty band, propagated from the posterior coefficient variances through the same nonlinear forward integration used for the point estimate, contains the vast majority of the interpolated trajectory across the full observation window (Figure 4b). None of the competing methods captures the observed oscillatory dynamics, likely due to the high collinearity and active Hill influence. SR3’s identified Hill coefficient is roughly one order of magnitude below RI-SINDy’s, and Nullcline-SINDy struggles to capture an accurate period of oscillation, drifting out of phase over the observation window.

**Table 3:** Hes1 mRNA-protein network trajectory error against the interpolated reference.

| Variable | Method | RMSE | Rel. $\ell_2$ |
| --- | --- | --- | --- |
| mRNA | RI-SINDy | **0.8964** | **0.3553** |
|  | SR3 | 4.0548 | 1.6074 |
|  | Nullcline-SINDy | 1.5225 | 0.6035 |
|  | Trad. SINDy | 1.1961 | 0.4741 |
| Protein | RI-SINDy | **1.2573** | **0.3308** |
|  | SR3 | 2.9015 | 0.7634 |
|  | Nullcline-SINDy | 2.8023 | 0.7373 |
|  | Trad. SINDy | 1.8189 | 0.4785 |

### 3.3 RI-SINDy in Noisy Synthetic Oscillators

Synthetic datasets were generated for two well known biological oscillators, a Goodwin limit cycle attractor and the NF-*κ*B regulatory network, to evaluate RI-SINDy against SR3, Nullcline-SINDy and Traditional SINDy under controlled noise. Both systems span three state variables and contain several nearly collinear candidate functions, but differ in the severity of that collinearity, as measured by pairwise correlation (Figures 5a and 7a).

**Figure 5:**
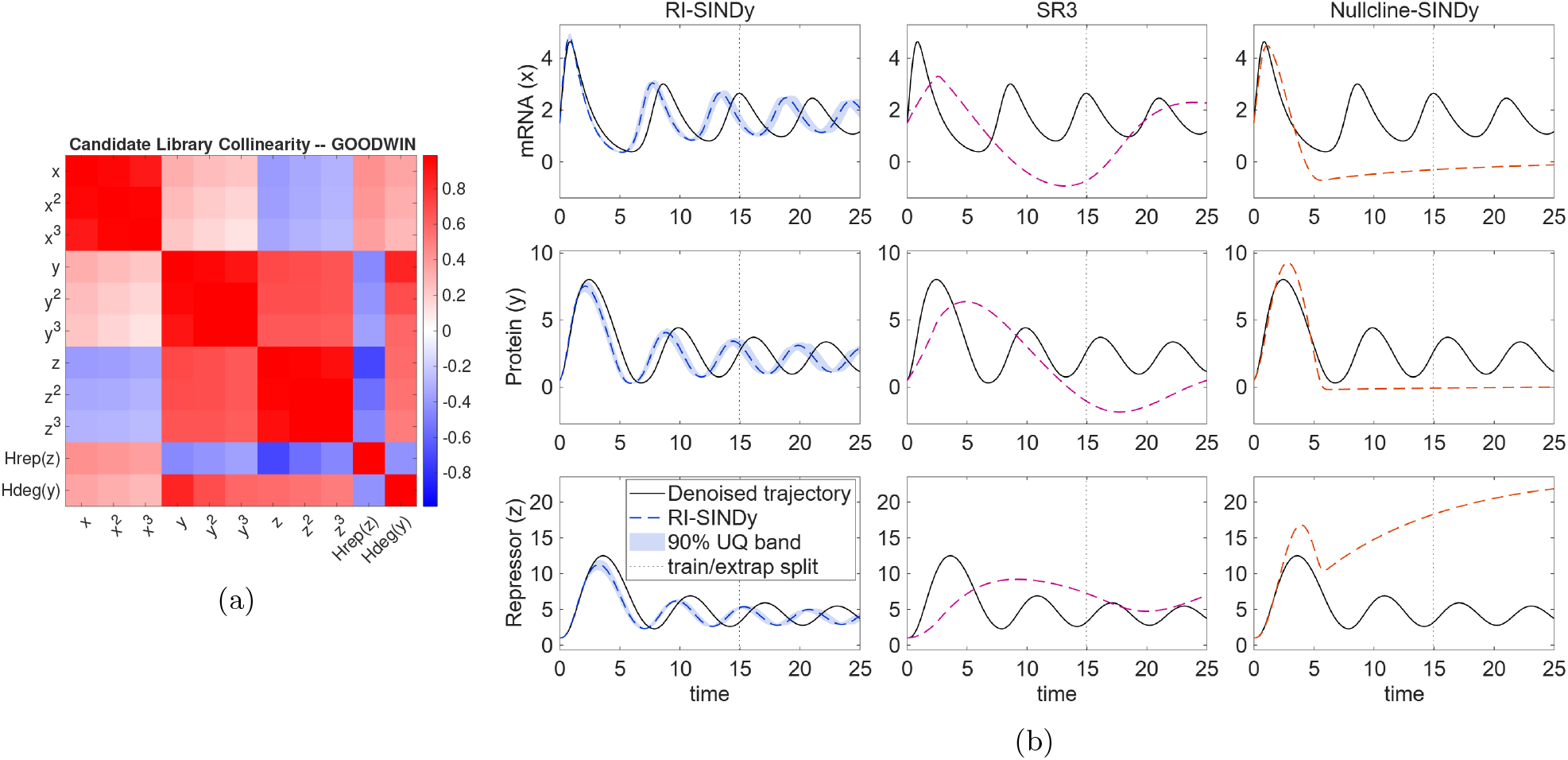
Candidate Library Collinearity and System Trajectory for the Goodwin System (a) Pairwise correlation of all candidate library terms for the Goodwin system. (b) Identified trajectories against the denoised control trajectory (black). RI-SINDy’s column includes its 90% uncertainty range.

#### 3.3.1 Goodwin Oscillator

The Goodwin attractor is a canonical three variable nonlinear ODE system describing a negative feedback loop in which a gene produces a repressive protein that pushes the system into a sustained oscillation of mRNA, protein and repressor concentrations [14, 43]. The model used to generate the data incorporates two distinct Hill-type nonlinearities, a Hill-type repression term in the mRNA equation and a saturating Michaelis-Menten degradation term in the protein equation [14], formulated as

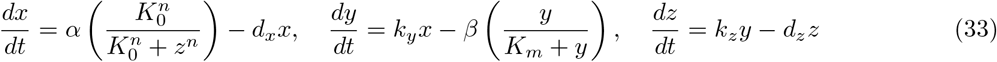

with control parameters *α* = 8, *d*_*x*_ = 0.6, *k*_*y*_ = 2, *β* = 5, *K*_*m*_ = 1, *k*_*z*_ = 1.5 and *d*_*z*_ = 0.8. 5% Gaussian noise was added to each data point, and every method was fit on the first three hundred of five hundred and one total time points, with the remaining two hundred reserved as an extrapolation test. Figure 5 shows the resulting candidate library correlations and the identified trajectories against the denoised control.

The mRNA, protein and repressor terms each form a highly correlated block with their respective polynomial powers, but crossed terms between the three state variables are mild, ranging from weak positive to weak negative correlation (Figure 5a).

RI-SINDy recovers the correct terms in every equation. Active coefficients are within 6% of their true values in the protein and repressor equations. The largest errors are in the mRNA equation (*x*: *−*0.7518 compared to true value *−* 0.6000, 25.3%; *H*_rep_(*z*): 9.3623 compared to true value 8.0000, 17.0%). SR3 spreads small nonzero weight across almost every candidate term in each equation. Nullcline-SINDy recovers both protein-equation terms with a mean error of 0.54%, but omits the *x* term in the mRNA equation and the *z* term in the repressor equation, underestimates *H*_rep_(*z*) and *y* by 37.6% and 25.7%, and adds spurious terms in both equations.

RI-SINDy achieves the lowest trajectory error of any method on every variable (Table 5), consistent with its coefficient vector matching all six active and all inactive terms in the control model to within roughly 25% (Table 4). SR3 returns roughly double RI-SINDy’s relative error. Nullcline-SINDy’s relative *l*_2_ errors of 0.94, 0.66 and 2.03 show that accurate coefficients in one equation do not yield accurate trajectories once the coupled equations are misidentified, with the repressor error exceeding the norm of the signal.

**Table 4:** Synthetically generated Goodwin model coefficients. Highlighted rows indicate terms present in the control model. SR3 discovered *K*_0_ = 3.42, *n* = 20.00 (hit its upper search bound), *K*_*m*_ = 3.25 against true values of *K*_0_ = 2, *n* = 3, *K*_*m*_ = 1.

| Term | Control |  |  | RI-SINDy |  |  | SR3 |  |  | Nullcline-SINDy |  |  |
| --- | --- | --- | --- | --- | --- | --- | --- | --- | --- | --- | --- | --- |
|  | mRNA | Prot. | Rep. | mRNA | Prot. | Rep. | mRNA | Prot. | Rep. | mRNA | Prot. | Rep. |
| Constant | 0 | 0 | 0 | 0 | 0 | 0 | 0.0413 | -0.1184 | -0.3110 | 0 | 0 | 0 |
| $x$ | -0.6000 | 2.0000 | 0 | -0.7518 | 2.0230 | 0 | 0.0003 | 0.1833 | 0.1911 | 0 | 1.9875 | 1.7615 |
| $y$ | 0 | 0 | 1.5000 | 0 | 0 | 1.4857 | -0.0563 | -0.0015 | 0.0783 | -0.1923 | 0 | 1.1145 |
| $z$ | 0 | 0 | -0.8000 | 0 | 0 | -0.7906 | -0.0007 | -0.0627 | -0.0039 | 0 | 0 | 0 |
| $H_{\text{rep}}(z)$ | 8.0000 | 0 | 0 | 9.3623 | 0 | 0 | 0.9802 | 1.0508 | -0.1709 | 4.9912 | 0 | 0 |
| $H_{\text{deg}}(y)$ | 0 | -5.0000 | 0 | 0 | -5.2834 | 0 | -0.3357 | -0.2702 | 0.1653 | 0 | -4.9772 | -10.0328 |

**Table 5:** Goodwin oscillator trajectory error against the noise-free control and coefficient recovery accuracy per state variable. Spurious terms, missing terms, and coefficient error use a |*ξ*| *<* 0.05 threshold for considering an estimated coefficient as effectively zero. Coefficient RMSE and mean relative error are calculated only over terms correctly identified as active for that variable.

| Variable | Method | RMSE | Rel. $\ell_2$ | Spurious | Missing | Coeff. RMSE | Mean Rel. Error |
| --- | --- | --- | --- | --- | --- | --- | --- |
| mRNA | RI-SINDy | **0.8169** | **0.4214** | **0** | **0** | **0.9693** | **21.16%** |
|  | SR3 | 1.6943 | 0.8741 | 2 | 1 | 7.0198 | 87.75% |
|  | Nullcline-SINDy | 1.8264 | 0.9423 | 1 | 1 | 3.0088 | 37.61% |
| Protein | RI-SINDy | **1.4081** | **0.4022** | **0** | **0** | 0.2011 | 3.41% |
|  | SR3 | 3.1666 | 0.9045 | 4 | 0 | 3.5827 | 92.72% |
|  | Nullcline-SINDy | 2.3175 | 0.6619 | **0** | **0** | **0.0184** | **0.54%** |
| Repressor | RI-SINDy | **1.6727** | **0.2799** | **0** | **0** | **0.0121** | **1.07%** |
|  | SR3 | 3.7633 | 0.6297 | 5 | 1 | 1.4217 | 94.78% |
|  | Nullcline-SINDy | 12.1340 | 2.0304 | 2 | 1 | 0.3855 | 25.70% |

#### 3.3.2 Noise Sensitivity

Figure 6 compares Traditional SINDy, E-SINDy and RI-SINDy on the Goodwin oscillator at noise levels of 0, 5, 10 and 15%. For each method and noise level, the sparsity parameter *λ* is chosen to balance correct term recovery against thresholding and prevent overfitting. RI-SINDy recovers all 6 active terms at every tested noise level, whereas Traditional SINDy recovers 4, 3, 1 and 2 of 6 and E-SINDy recovers 4, 2, 1 and 0 of 6. RI-SINDy also returns the lowest relative *L*_2_ error at every level: 0.200, 0.394, 0.274 and 0.675, when compared to 0.633–2.525 for the other methods. RI-SINDy retains 1, 3, 3 and 2 low magnitude spurious terms at the four levels and is the only method with a spurious term at 0% noise. Traditional SINDy identifies up to 6 spurious terms at 5% noise, while E-SINDy identifies at most 2; their low spurious counts at 15% show that they recover few terms at all.

**Figure 6:**
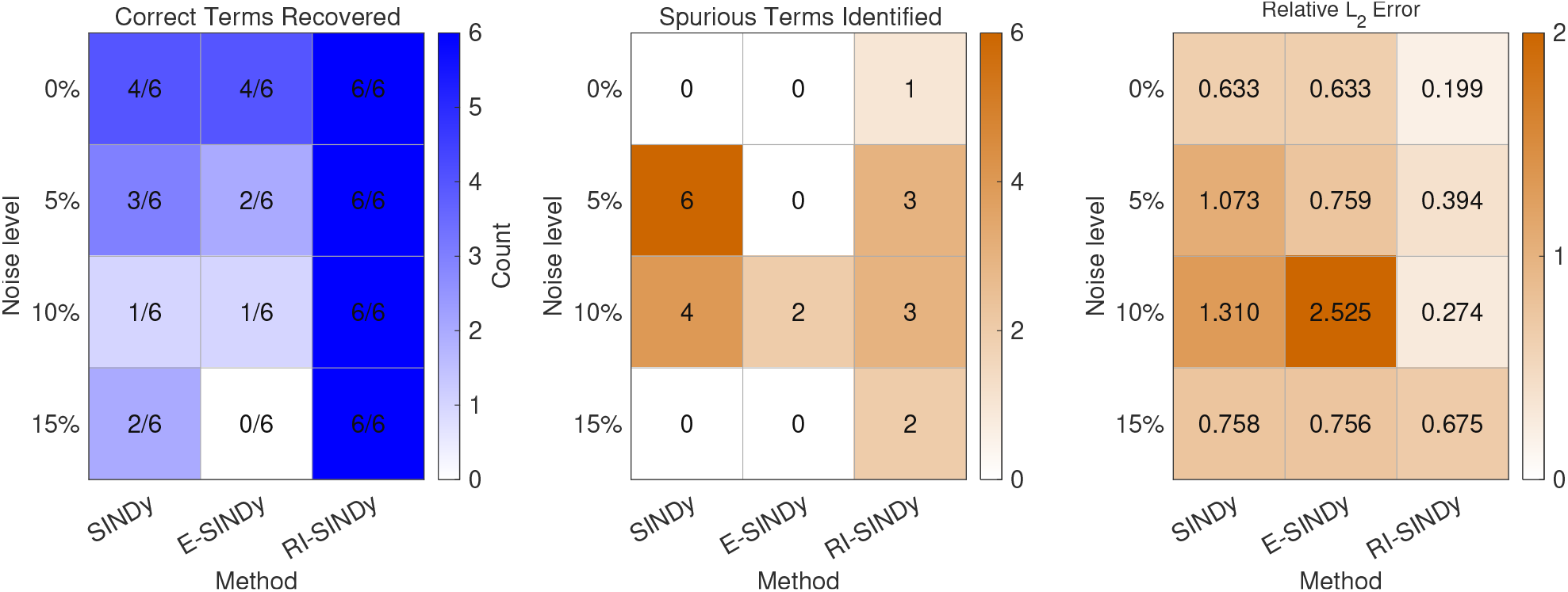
Noise sensitivity analysis on the Goodwin attractor for Traditional SINDy, E-SINDy and RI-SINDy. Correct terms is the number of recovered terms that are in the control model, Spurious terms are counted as any nonzero coefficient outside the control support, without the |*ξ*| *<* 0.05 cutoff used in Table 5.

#### 3.3.3 NF-*κ*B Network

NF-*κ*B is a transcription factor that plays a significant role in immune response and inflammation. NF-*κ*B oscillates in concentration between the cytoplasm and nucleus as a result of a negative feedback loop with its inhibitor I*κ*B, which is modulated by the kinase IKK in response to inflammatory signals [1, 6, 44]. This process can be captured in a three variable ODE system, consisting of a single Hill function capturing the NF-*κ*B driven transcription of IKK [6, 45], formulated as

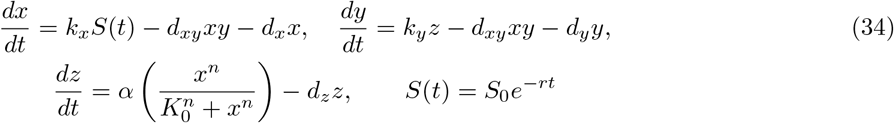

where *S*(*t*) is the external inflammatory signal driving NF-*κ*B synthesis, a pulse of magnitude *S*_0_ at *t* = 0 followed by exponential decay at rate *r*. 10% Gaussian noise was added to each data point, and no train and extrapolation split was applied for this system.

The NF-*κ*B dataset is a single pulse response, where each state rises transiently and then decays over the window, with no sustained oscillation (Fig. 7b), so it lies outside the limit-cycle regime that motivates this method. It satisfies the conditions of §2.4 only partially as the boundary displacement is small relative to the accumulated drift. Only the IKK equation uses the balance. By Eq.(19), the estimate of *ξ*_*R*_ in that equation carries a bias of *−*[*z*(*T*) *− z*(0)]*/*(*TD*_*R*_); here *z*(0) *≈* 0.6 and *z*(*T*) *≈* 0.05 at *T* = 50, giving a relative error of *ρ*_*z*_ = |*z*(*T*) *−z*(0) |*/*(*T ξ*_*R*_*D*_*R*_) *≈*7%. We generate data in the context of a transient system rather than an oscillatory system to inspect the coefficient recovery in systems that fail to perfectly fit the applicability conditions of RI-SINDy. These results should therefore be read as a test of drift balance on a decaying transient with a boundary term of this size, not as a limit cycle test.

**Figure 7:**
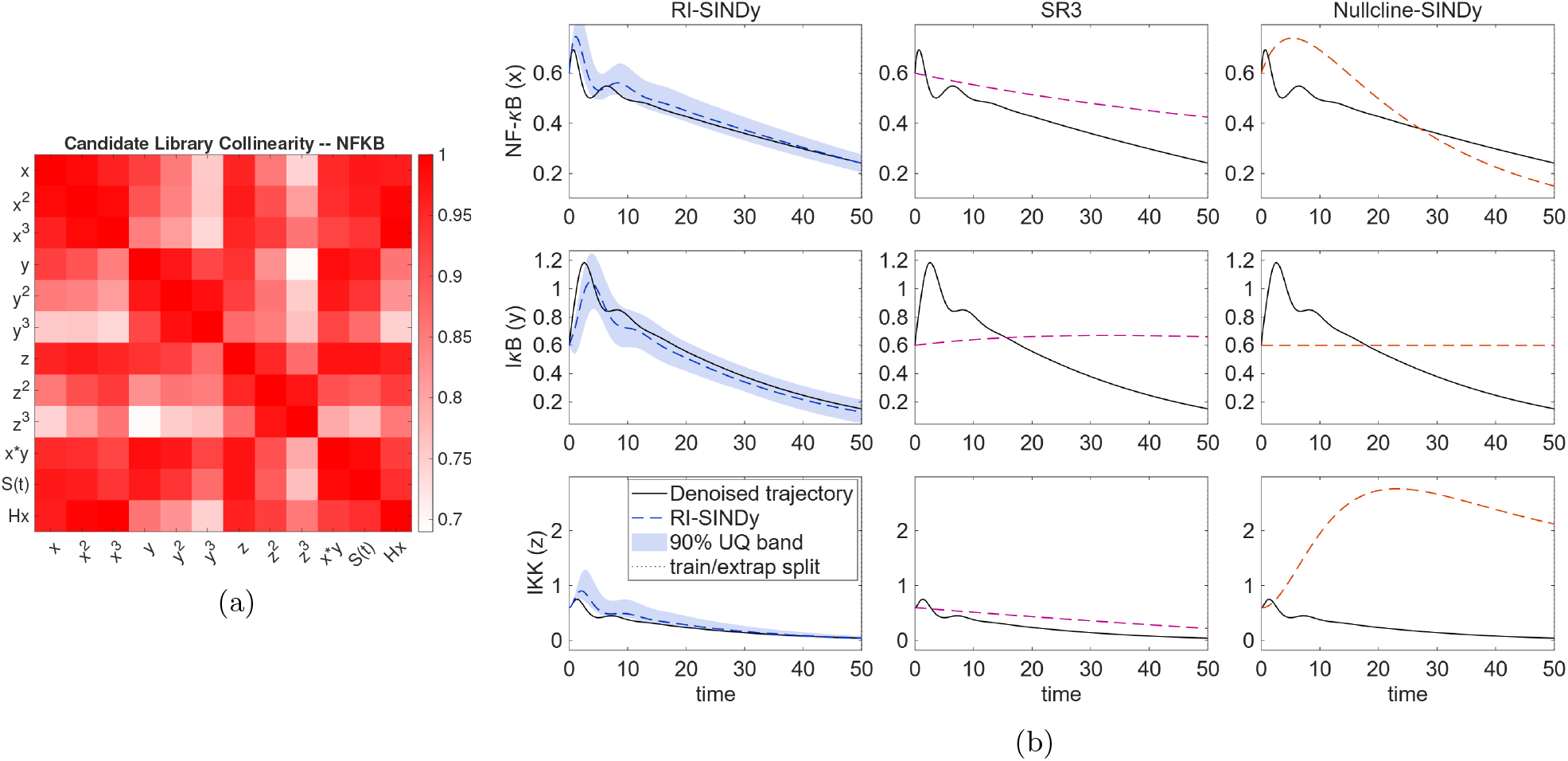
Candidate Library Collinearity and System Trajectory for the NF-*κ*B System (a) Pairwise correlation of all candidate library terms for the NF-*κ*B system. The colorbar here spans only about 0.7 to 1.0, since no pair of terms in this library falls below that range. (b) Identified trajectories against the denoised control trajectory (black). RI-SINDy’s column includes its 90% uncertainty range.

Figure 7 shows the corresponding correlation structure and identified trajectories for this system.

Every visible cell in Figure 7a is strongly correlated. The range of collinearity spans only from roughly 0.7 to 1.0, meaning the weakest correlation present anywhere in the thirteen term library is still stronger than almost every correlation observed in the Goodwin library.

The largest coefficient errors occur in the I*κ*B equation (Table 6), where all three active terms are underestimated in magnitude: *z* (0.9731 vs. 1.8000, 45.9%), *y* (*−* 0.2052 vs. *−* 0.3000, 31.6%) and *x· y* (*−* 0.8421 vs. *−* 1.2000, 29.8%). Under all three priors, the largest posterior standard deviations in this equation fall on these same terms (Table 8; Spike-and-Slab: *z* 0.1598, *x · y* 0.1468, *y* 0.0702), and as such, the uncertainty estimates identify where the fit is least reliable. SR3 returns coefficients of order 10^*−*2^ or smaller for every term, and Nullcline-SINDy underestimates the NF-*κ*B-equation coefficients by 33–63% and returns no terms in the I*κ*B equation.

**Table 6:** Synthetically generated NF-*κ*B model coefficients. Highlighted rows indicate terms present in the control model. SR3 discovered *K*_0_ = 0.50, *n* = 1.00 against true values of *K*_0_ = 1.5, *n* = 3.

| Term | Control |  |  | RI-SINDy |  |  | SR3 |  |  | Nullcline-SINDy |  |  |
| --- | --- | --- | --- | --- | --- | --- | --- | --- | --- | --- | --- | --- |
| | $NF-\kappa B$ | $I\kappa B$ | $IKK$ | $NF-\kappa B$ | $I\kappa B$ | $IKK$ | $NF-\kappa B$ | $I\kappa B$ | $IKK$ | $NF-\kappa B$ | $I\kappa B$ | $IKK$ |
| Constant | 0 | 0 | 0 | 0 | 0 | 0 | -0.0002 | -0.0006 | -0.0002 | 0 | 0 | 0 |
| $x$ | -0.4000 | 0 | 0 | -0.4241 | 0 | 0 | -0.0005 | -0.0007 | -0.0004 | 0 | 0 | 0 |
| $x^2$ | 0 | 0 | 0 | 0 | 0 | 0 | -0.0010 | 0.0004 | -0.0005 | 0 | 0 | 0 |
| $x^3$ | 0 | 0 | 0 | 0 | 0 | 0 | -0.0016 | 0.0052 | 0.0001 | 0 | 0 | 0 |
| $y$ | 0 | -0.3000 | 0 | 0 | -0.2052 | 0 | -0.0005 | -0.0006 | -0.0007 | 0 | 0 | 0 |
| $y^2$ | 0 | 0 | 0 | 0 | 0 | 0 | -0.0007 | -0.0005 | -0.0013 | 0 | 0 | 0 |
| $y^3$ | 0 | 0 | 0 | 0 | 0 | 0 | -0.0009 | -0.0003 | -0.0017 | 0 | 0 | 0 |
| $z$ | 0 | 1.8000 | -0.8000 | 0 | 0.9731 | -0.6881 | -0.0008 | 0.0007 | -0.0008 | 0 | 0 | 0 |
| $z^2$ | 0 | 0 | 0 | 0 | 0 | 0 | -0.0020 | 0.0067 | -0.0012 | 0 | 0 | 0 |
| $z^3$ | 0 | 0 | 0 | 0 | 0 | 0 | -0.0039 | 0.0169 | -0.0011 | 0 | 0 | 0 |
| $x \cdot y$ | -1.2000 | -1.2000 | 0 | -1.0803 | -0.8421 | 0 | -0.0009 | -0.0002 | -0.0012 | -0.3445 | 0 | 0 |
| $S(t)$ | 1.0000 | 0 | 0 | 0.9581 | 0 | 0 | -0.0004 | -0.0003 | -0.0005 | 0.1922 | 0 | -0.1960 |
| $H(x)$ | 0 | 0 | 8.0000 | 0 | 0 | 6.9522 | -0.0005 | -0.0010 | -0.0088 | 0 | 0 | 3.0503 |

RI-SINDy outperforms the other tested methods by a wide margin on all three variables (Table 7), with relative *l*_2_ errors of 0.1139 on I*κ*B and 0.2573 on IKK. Nullcline-SINDy’s relative *l*_2_ error on IKK exceeds one, indicating an error larger than the signal itself and evidence that the model carries no useful predictive information about that state variable under this noise level. Figure 7b confirms this, with RI-SINDy tracking the qualitative decay of all three state variables including the initial transient spike.

**Table 7:** NF-*κ*B network trajectory error against the noise-free control and coefficient recovery accuracy per state variable. Spurious terms, missing terms, and coefficient error use a |*ξ* |*<* 0.05 threshold for considering an estimated coefficient as effectively zero. Coefficient RMSE and mean relative error are calculated only over terms correctly identified as active for that variable. If global sparsity constraints force a particular variable to contain no terms, its coefficient statistics are marked by a ‘–’ metric.

| Variable | Method | RMSE | Rel. $\ell_2$ | Spurious | Missing | Coeff. RMSE | Mean Rel. Error |
| --- | --- | --- | --- | --- | --- | --- | --- |
| NF-κB | RI-SINDy | **0.0311** | **0.0751** | **0** | **0** | **0.0745** | **6.73%** |
|  | SR3 | 0.1147 | 0.2772 | 0 | 3 | — | — |
|  | Nullcline-SINDy | 0.1118 | 0.2700 | 0 | 1 | 0.8320 | 76.04% |
| IκB | RI-SINDy | **0.0664** | **0.1139** | **0** | **0** | **0.5231** | **35.79%** |
|  | SR3 | 0.3222 | 0.5529 | 0 | 3 | — | — |
|  | Nullcline-SINDy | 0.2864 | 0.4914 | 0 | 3 | — | — |
| IKK | RI-SINDy | **0.0747** | **0.2573** | **0** | **0** | **0.7451** | **13.55%** |
|  | SR3 | 0.1823 | 0.6279 | 0 | 2 | — | — |
|  | Nullcline-SINDy | 2.1177 | 7.2923 | 1 | 1 | 4.9497 | 61.87% |

**Table 8:** RI-SINDy identified posterior standard deviations for dimension *j* = 2 of the NF-*κ*B model under three statistical priors, Spike-and-Slab (SS), Laplace (LAP) and Horseshoe (H). These deviations correspond to the I*κ*B equation.

| **Term** | **SS** | **LAP** | **$H$** |
| --- | --- | --- | --- |
| 1 | 0.0107 | 0.0212 | 0.0018 |
| $S$ | 0.0097 | 0.0207 | 0.0011 |
| $x$ | 0.0209 | 0.0439 | 0.0024 |
| $x^2$ | 0.0160 | 0.0326 | 0.0018 |
| $x^3$ | 0.0295 | 0.0172 | 0.0044 |
| $y$ | **0.0702** | **0.0627** | **0.0095** |
| $y^2$ | 0.0024 | 0.0052 | 0.0003 |
| $y^3$ | 0.0006 | 0.0013 | 0.0001 |
| $z$ | **0.1598** | **0.1188** | **0.0322** |
| $z^2$ | 0.0356 | 0.0087 | 0.0055 |
| $z^3$ | 0.0011 | 0.0024 | 0.0001 |
| $x \cdot y$ | **0.1468** | **0.1725** | **0.0330** |
| $H(x)$ | 0.0000 | 0.0000 | 0.0000 |

## 4 Discussion

This work introduced RI-SINDy, a framework designed for biological systems containing known saturating regulatory nonlinearities. Rather than estimating these terms within the shared sparse regression, RI-SINDy separates them from the candidate library and estimates their coefficients through a drift-balance condition based on the long term behavior of bounded trajectories. This decomposition preserves the mechanistic interpretation of regulatory kinetics while removing the regulatory term’s collinearity from the standard basis terms in the regression, and is compatible with the constrained regression and uncertainty quantification machinery already established in the SINDy literature.

Across the three systems tested, RI-SINDy’s advantage scaled with the severity of library collinearity. On the Hes1 network, fit to real experimental measurements, RI-SINDy achieved the lowest trajectory error of any method on both state variables, while other methods struggled to capture the observed oscillation. On the synthetic Goodwin oscillator, RI-SINDy recovered a sparse model structurally aligned with the six term control system and achieved the lowest trajectory error of any tested method, while SR3 spread weight across nearly every candidate term and Nullcline-SINDy omitted active terms in the mRNA and repressor equations and produced trajectories that failed to track the oscillation. On the more severely collinear NF-*κ*B network, RI-SINDy again recovered the most kinetically plausible model and the lowest error on all three state variables, but also produced one clear coefficient error, undershooting the magnitude of the *z* production term in the I*κ*B equation by about a factor of two. Its own uncertainty quantification flagged this term directly, with one of the largest posterior standard deviations of any term in that equation.

The results suggest that the conditioning benefit arises from removing the regulatory columns from the matrix that is inverted (§2.2, Fig. 2). While regularization and decorrelation methods can mitigate conditioning for individual libraries, their effectiveness becomes increasingly limited as additional Hill functions introduce further collinearity. Once these terms are estimated independently through the drift-balance condition, the established regression and uncertainty quantification methodologies can operate on a better conditioned system.

The conditions for where RI-SINDy is applicable include each state entering a drift balance that remains bounded with net displacement small relative to its accumulated drift, the regulatory term’s mean over the window being bounded away from zero, and the functional form of the regulatory mechanism being known (§2.4). Most inferences generated for these results are from a single noise level per condition, and the Hes1 hyperparameters were selected from literature regarding Hes1 2-state dynamical models [1].

Several directions follow directly from these limitations. Relaxing the single term restriction to accommodate multiple coupled regulatory nonlinearities, most likely through an additional balance condition or an independent source of constraint, is the most immediate priority. Jointly identifying Hill hyperparameters such as cooperativity and half-saturation alongside the regulatory coefficient, rather than fixing them in advance, would reduce the method’s reliance on external optimizations to discover hyperparameters. A formal convergence analysis of the alternating procedure could be performed to address the remaining numerical limitations of finite length drift-balance calculations. Though the proposed drift-balance condition is introduced per state dimension, implying a resistance to higher dimensional regressions, testing RI-SINDy on more complex networks and stochastic formulations would directly quantify how far the drift-balance approach generalizes beyond the low dimensional systems considered here.

This work suggests that supplementing a sparse regression with an independent source of physical information, rather than continuing to regularize or stabilize a poorly conditioned term within it, can improve model discovery without losing interpretability. While developed for biological regulatory systems, the same principle may prove useful whenever a small number of known nonlinear mechanisms dominate the conditioning of an otherwise sparse identification problem.

## Ethics Statement

This study did not involve human participants, human tissue, or animal experiments. The Hes1 data were obtained from previously published measurements [9]; all other data were generated from established mathematical models.

## Data Availability

The MATLAB implementation of RI-SINDy, along with the scripts used to generate the synthetic Goodwin and NF-*κ*B datasets and reproduce the figures and tables in this work, is available at https://github.com/NicholasJLucenti/RI-SINDy.

## Author Contributions

Conceptualization: N.L., M.B.Y.; Methodology: N.L.; Literature Search and Screening: N.L. & M.B.Y.; Data Extraction: M.B.Y. Writing the Original Draft: N.L.; Review & Editing: M.B.Y.; Supervision: M.B.Y.

